# Sequential digestion with Trypsin and Elastase in cross-linking/mass spectrometry

**DOI:** 10.1101/450981

**Authors:** Therese Dau, Kapil Gupta, Imre Berger, Juri Rappsilber

## Abstract

Cross-linking/mass spectrometry has become an important approach for studying protein structures and protein-protein interactions. The amino acid composition of some protein regions impedes the detection of cross-linked residues, although it would yield invaluable information for protein modelling. Here, we report on a sequential digestion strategy with trypsin and elastase to penetrate regions with a low density of trypsin cleavage sites. We exploited intrinsic substrate recognition properties of elastase to specifically target larger tryptic peptides. Our application of this protocol to the TAF4-12 complex allowed us to identify cross-links in previously inaccessible regions.

---

Trypsin is the enzyme of choice in mass spectrometry (MS)-based proteomics. The favorable behavior of tryptic peptides during mass spectrometric analysis is one of the main reasons for this^1,2^. However, as trypsin cleaves after Arg and Lys, peptides generated in protein regions with a low Arg/Lys density are very long, making them potentially non-observable by mass spectrometry and thus resulting in poor coverage of these regions.

This problem is particularly relevant for cross-linking/mass spectrometry (CLMS). In CLMS, protein proximities and conformations are preserved through the introduction of covalent bonds. The detection of these bonds as cross-linked peptide pairs by MS is translated into distance constraints^3–5^. Currently, the most frequently used cross-linkers are Nhydroxysuccinimide (NHS) esters that primarily target the ∊-amino groups of Lys residues, and to a lesser extent also react with the hydroxyl groups of Ser, Thr and Tyr^6,7^. CLMS critically depends on good sequence coverage to reveal structural information for the whole protein. Cross-linking naturally involves two peptides, which aggravates the problem of overly large peptides. This is exacerbated when the cross-linker reacts with two lysine residues as two potential cleavage sides for trypsin are removed. The stabilization of proteins through cross-linking and the destruction of potential trypsin cleavage sites may further affect the efficiency of trypsin digestion and lead to an increase in missed cleavages.

Alternative proteases to trypsin target basic (LysC, ArgC), acidic (AspN and GluC) or aromatic (chymotrypsin) amino acids^8,9^. Although usage of these proteases improves sequence coverage^10^ and has been used in parallel to trypsin to enhance identification of cross-linked residues^11,12^, some protein regions such as the N-terminal region of TAF4, a subunit of transcriptional factor II D, still cannot be accessed (Figure 1). The existence of TAF4b, a cell type-specific TAF4 paralog, with a unique N-terminal region, points to important functions and interactions mediated through the N-terminal region of TAF4^13^. Cross-linked residues in this region would therefore provide important information on the function and structure of TAF4.

**Figure 1.**
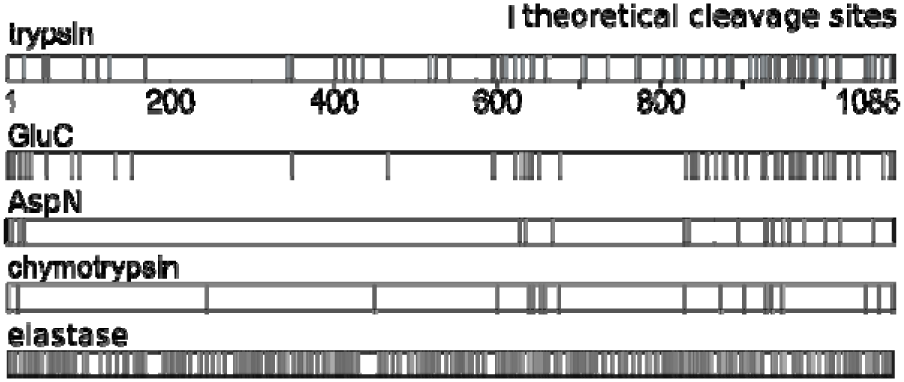
Theoretical cleavage sites in TAF4.

Proteases with broader specificity like elastase, proteinase K and pepsin have the potential of complementing more selective proteases and indeed have been used in membrane proteomics^14,15^ to detect phosphorylation sites^16^, and even in CLMS^17^, with varying levels of success. Elastase, for example, cleaves after Ala, Val, Leu, Ile, Ser and Thr^14^ and should be able to cleave the N-terminal region of TAF4 at many sites (Figure 1). This broad specificity leads to two potential problems. If cleavage should be complete, the resulting peptides would be very small. However, cleavage of elastase is not expected to be complete. Analyzes of elastase efficiency on peptides with varying length revealed elastase to have an extended substrate binding region^18,19^. Short peptides that do not cover the complete substrate-binding region are less efficiently cleaved by elastase. While this addresses the potential problem of peptide length, it leads to another problem. Having many missed cleavage sites leads elastase to generate very complex mixtures with often overlapping peptides^14^. This problem is exacerbated in cross-linked peptides as two peptides are involved, resulting in a combinatorial increase of complexity. Recent work by our lab revealed a strong substrate preference of proteases for longer peptides^12^. As a consequence, in a sequential digest, preferentially long peptides are cleaved by the second protease while short peptides remain undigested. The complexity of an elastase digest might therefore be vastly reduced by first using trypsin. Only long tryptic peptides should be good substrates to elastase while any sequence range of a protein that is covered by short tryptic peptides would remain unaffected by elastase treatment.

In this study, we investigate the hypothesis that using multiple enzymes positively influences the detection odds of peptides. Sequential digestion influences the digestion outcome and thus is more than the sum of the individual digestion approaches. As a result, we detect cross-links in the difficult to digest N-terminal region of TAF4.

## Material and methods

### Sample preparation

Human TAF4-TAF12 complex was co-expressed in Sf21 insect cells using the MultiBac system^20^. DNA encoding for a N-terminal hexa-histidine tag and a protease cleavage site for tobacco etch virus (TEV) NIa protease was added to the 5’ end of the TAF4 open reading frame and cloned into pFL plasmid along with TAF12. Baculoviurus generation, cell culture infection and protein production was carried out following published protocols^21^. Cell pellets were resuspended in Talon Buffer A (25 mM Tris pH 8.0, 150 mM NaCl, 5 mM imidazole with complete protease inhibitor (Roche Molecular Biochemicals)). Cells were lysed by freeze-thawing (twice), followed by centrifugation at 40,000 g in Ti70 rotor for 60 min to clear the lysate. The TAF4-TAF12 complex was first bound to talon resin, pre-equilibrated with Talon Buffer A, followed by washes with Talon Buffer A, then Talon Buffer HS (25 mM Tris pH 8.0, 1 M NaCl, 5 mM imidazole and complete protease inhibitor) and then again with Talon Buffer A. The TAF4-TAF12 complex was eluted using Talon Buffer B (25 mM Tris pH 8.0, 150 mM NaCl, 200 mM imidazole and complete protease inhibitor). Fractions containing the TAF4-TAF12 complex were dialyzed overnight against MonoQ Buffer A (25 mM HEPES pH 7.5, 100 mM NaCl, 1 mM DTT and complete protease inhibitor). The complex was further purified using ion exchange chromatography (IEX) with a MonoQ column pre-equilibrated with MonoQ Buffer A. After binding, the column was washed with MonoQ Buffer A and TAF4-TAF12 was eluted using a continuous gradient of MonoQ Buffer B (25 mM HEPES pH 7.5, 1000 mM NaCl, 1 mM DTT and complete protease inhibitor) from 0-100% linear gradient. The complex was further purified by size exclusion chromatography (SEC) with a SuperoseS6 10/300 column in SEC buffer (25 mM Tris pH 8.0, 150 mM NaCl, 1 mM DTT and complete protease inhibitor).

TAF4-TAF12 (1.5 µg per condition) complexes were cross-linked with bis(sulfosuccinimidyl)suberate (BS3) (Thermo Fisher Scientific, Rockford, IL) at weight-to-weight ratio of 1:1 and incubated on ice for 2 hours. The mixture was incubated on ice for 1 hour after saturated bicarbonate (50 molar excess) was added to quench the reaction. Frozen *S. pombe* cells were ground and 1g of yeast powder was resuspended in 2 ml RIPA (Sigma-Aldrich, St. Louis, MO) supplemented with cOmplete™ (Roche, Basel). Cell debris was moved via centrifugation at 1200 g for 15 minutes. All samples were separated by SDS–PAGE on a 4-12% Bis Tris gel (Life Technologies, Carlsbad, CA) and stained using Imperial Protein Stain (Thermo Fisher Scientific, Rockford, IL). Appropriate bands were cut and proteins were first reduced with DTT and then alkylated with iodoacetamide. Samples were incubated with trypsin (13 ng µl^−1^) (Pierce™, Thermo Fisher Scientific, Rockford, IL) or elastase (15 ng µl^−1^) (Promega) respectively at 37°C for 16 hours. For the sequential digestion, elastase (15 ng µl^−1^) was added to the trypsin digest and incubated at 37°C for 30 minutes. Following the digestion, peptides were purified on C18 StageTips using standardized protocols^22^.

### LC-MS/MS

TAF4-TAF12 complexes were analyzed on an Orbitrap Fusion™ Lumos™ Tribrid™ (Thermo Fisher Scientific, Rockford, IL) and yeast lysates were analyzed on an Orbitrap Elite (Thermo Fisher Scientific, Rockford, IL). Both were coupled online to an Ultimate 3000 RSLCnano Systems (Dionex, Thermo Fisher Scientific, UK). Peptides were loaded onto an EASY-Spray™ LC Column (Thermo Fisher Scientific, Rockford, IL) at a flow rate of 0.300 μL min^−1^ using 98% mobile phase A (0.1% formic acid) and 2% mobile phase B (80% acetonitrile in 0.1% formic acid). To elute the peptides, the percentage of mobile phase B was first increased to 40% over a time course of 110 minutes followed by a linear increase to 95% in 11 minutes.

Full MS scans for yeast lysates were recorded in the orbitrap at 120,000 resolution with a scan range of 300-1700 m/z. The 20 most intense ions (precursor charge ≥2) were selected for fragmentation by collision-induced disassociation and MS2 spectra were recorded in the ion trap (2.0E04 ions as a minimal required signal, 35 normalized collision energy, dynamic exclusion for 40 s). Both MS1 and MS2 were recorded in the orbitrap for TAF4-TAF12 complexes (mass resolution 120,000, scan range 300-1700 m/z, dynamic exclusion for 60 s, precursor charge ≥3). Higher-energy collision dissociation was used to fragment peptides (30% collision energy, 5.0E04 AGC target, 60 ms maximum injection time).

### Data analysis

MaxQuant software^23^ (version 1.5.2.8) employing the Andromeda search engine^24^ was used to analyze the whole cell lysates of *S. Pombe*. We used the PombeBase database^25^ with carbamidomethylation of cysteine as a fixed modification and oxidation of methionine as a variable modification. MS accuracy was set to 4.5 ppm and MS/MS tolerance to 20 ppm. For trypsin, we allowed up to 2 miscleavages. Specificity of elastase was defined as cleavage after Ala, Val, Leu, Ile, Ser and Thr but not before Pro. Specificity for the trypsin and elastase digest was set to cleavage after Ala, Val, Leu, Ile, Ser, Thr, Arg and Lys, except if it was followed by Pro. For all digests containing elastase, up to 10 miscleavages were allowed. The peaklist for identification of cross-linked peptides was generated using MS convert (ProteoWizard) for tryptic digests with the following settings: peakPicking 2-; msLevel 2-; MS2Denoise 20,100, false or Maxquant version 1.6.2.3 with default setting except increasing the “Top MS/MS peaks per 100 Da” to 100 for elastase containing digests. Resulting mgf or apl files were searched using Xi software^12,26^ version: 1.6.731 with MS accuracy set to 4 ppm and MS/MS tolerance to 20 ppm. The only fixed modification was the carbamidomethylation of cysteine. Variable modifications were oxidation of methionine, amidated and hydrolyzed BS3. The cross-linker was BS3 with Lys, Ser, Thr and Tyr as the only cross-linkable residues. Enzyme specificities were the same as described for Mascot. Four miscleavages were allowed for the trypsin digest and 11 miscleavages were allowed for the other digests. An additional 2 missing isotopes was allowed in the custom settings. False discovery rates (FDR) were estimated using xiFDR^27^ version 1.1.26.58 with 5 % FDR at peptide-pair level for the TAF4-12 complex and 5 % FDR at peptide-pair and link-level. Linear peptides of the cross-linked TAF4-12 complex were identified using Mascot software^28^ with carbamidomethylation of cysteine as a fixed modification and the following variable modifications: oxidation of methionine, hydrolyzed BS3 (mass: 156.0786 Da) and amidated BS3 (mass: 155.0946 Da). Chromatograms of MS1 were extracted and peak areas quantified using Skyline ^29^.

### In-silico digest

**The** TAF4-TAF12 complex was digested in-silico using trypsin, elastase or a combination of trypsin and elastase. We assumed a complete digestion with no miscleavages for the in-silico digestion and a minimum length of 5 amino acids.

## Results and discussion

Four different types of digestion – trypsin, elastase, sequential trypsin-elastase and elastase-trypsin - were applied to a cross-linked TAF4-TAF12 complex. We compared these digests to assess the influence of substrate length on the cleavage behavior of elastase. The peptide length of observed tryptic peptides without any modifications ranged from 5 to 65 amino acids, which is very similar to the theoretical peptide length distribution (Figure 1a). However, the median of peptides digested by elastase, trypsin-elastase or elastase-trypsin was considerably higher (12 amino acids) than the predicted median of 6 amino acids (Figure 1b - d). This is in line with previous studies on short peptides reporting that the activity of elastase is dependent on the substrate length N-terminal of the cleavage site^18,19^. Our results suggest that a similar restriction might also apply to longer peptides and to the substrate length C-terminal from the cleavage site.

The majority of elastase-derived peptides are larger than five amino acids and the maximal length of 29 amino acids is considerably shorter than that of trypsin (65). Despite the favorable peptide length distribution of elastase-digested peptides, fewer peptide spectrum matches (546 ±74 PSM) were identified compared to peptides digested by trypsin alone (869 ±170 PSM), trypsin-elastase (1116 ±164 PSM) or elastase-trypsin (1111 ±100 PSM) (Figure 1e). One possible explanation for the reduced identification of elastase peptides could be the missing positive C-terminal charge that tryptic peptides have. To this end, we analyzed semi-tryptic peptides, as they should distribute equally between those featuring a tryptic C-terminus and a tryptic N-terminus. Semi-tryptic peptides with a tryptic C-terminus were identified more often (70 ±2%), thus suggesting that the positive C-terminal charge is beneficial for detection (Figure 1f). Another disadvantage of elastase is that frequently missed cleavage sites and overlapping peptides lead to complex peptide mixtures. By digesting proteins with trypsin first, the amount of available elastase cleavage sites might be reduced, as short tryptic peptides should be protected from elastase due to their size. To test this hypothesis, we quantified tryptic peptides before and after the addition of elastase. Peptides that had been reduced by more than half were labeled as “reduced”. The remaining peptides were categorized as “unchanged”. Peptides were divided into three groups depending on their length (≤ 12, 13-20 and >20). The number of reduced peptides, 19, 70 or 113 peptides, respectively, increased with peptide size (Figure 1g). The label-free quantification confirms that elastase digests preferentially long peptides, thus initial trypsin digestion reduces elastase-induced complexity. Indeed, analyzing the 404-420 region in TAF4 revealed that trypsin treatment protected this region from elastase digestion. Consequently, the complexity within the sequential trypsin-elastase digest was reduced compared to the digest from elastase alone or elastase-trypsin (Figure 2h).

**Figure 2.**
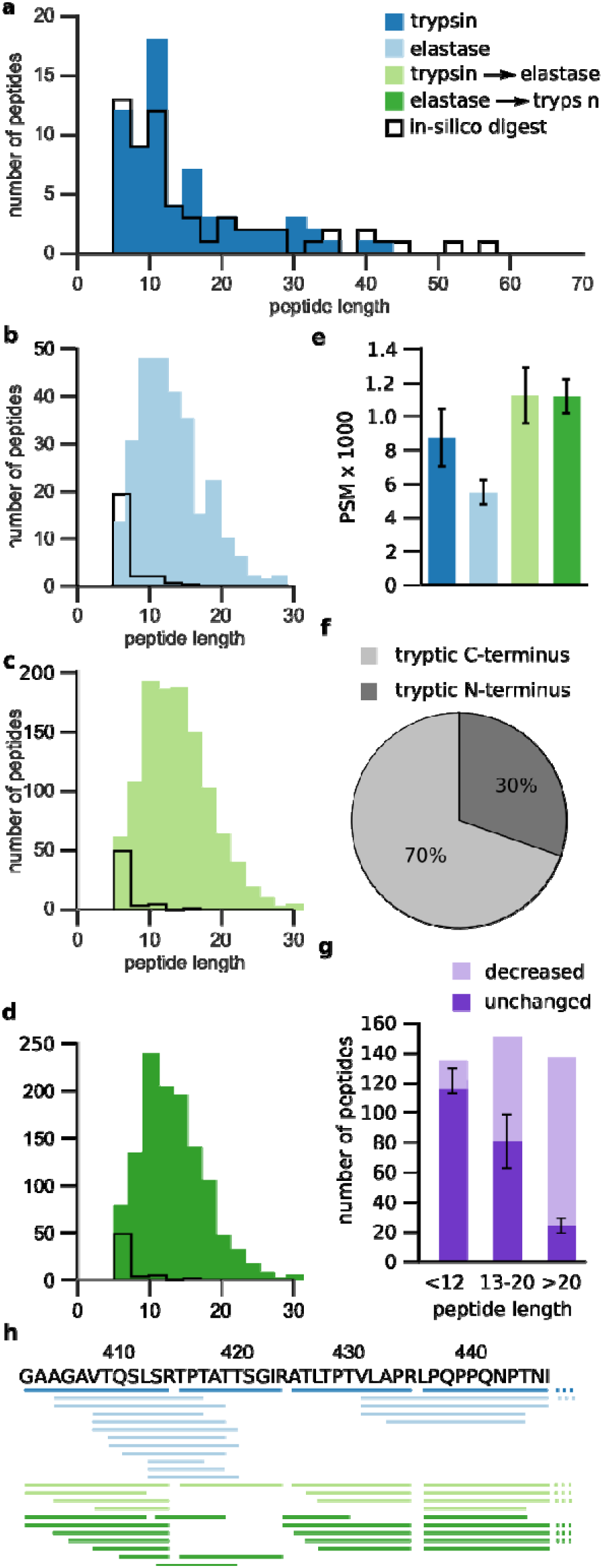
Impact of elastase when following trypsin in a sequential digestion of the TAF4-TAF12 complex. **(a-d)** Peptide length distribution of observed and theoretical peptides (full cleavage, no modifications) for trypsin (a), elastase (b), sequential trypsin-elastas (c) and elastase-trypsin **(d)** digestion. **(e)** Number of peptide-spectrum matches (PSM) for different proteases using Mascot. **(f)** Number of semi-tryptic peptides having tryptic C-terminus or tryptic N-terminus, respectively, in sequential trypsin-elastase digest. (**g**) Intensity change of tryptic peptides between tryptic digest and sequential trypsin-elastase digest using label-free quantitation in Skyline. The peptides were divided into three groups depending on their length (<12, 13-20 and >20 residues). Peptides with a 2-fold reduction or lower were labeled “decreased”. **(h)** TAF4 (residue 827-880) with the observed cleavage pattern of trypsin, elastase and the sequential trypsin-elastase treatment. Identified peptides are represented as lines below the protein sequence. Data shown are the mean ±standard deviation (SD) of three independent digestions. Trypsin (dark blue), elastase (light blue), sequential trypsin-elastase (light green) and sequential elastase-trypsin digest (dark green).

In the next step, we analyzed the effect of our sequential trypsin-elastase on the detection of cross-linked peptides. As a proof of principle, we first analyzed BS3 cross-linked human serum albumin (HSA). As expected, the increase in complexity and the missing C-terminal charge associated with elastase digestion adversely affected the identification of cross-linked peptides. While digestion of the BS3 cross-linked complexes with trypsin allowed us to identify 152 cross-links, digestion with elastase only led to the identification of 42 cross-links (Figure 3a). This situation improved when adding trypsin to the elastase digest. On one hand, adding trypsin to the elastase-digested peptides increased the complexity of the samples, but on the other hand it produced semi-tryptic and tryptic peptides featuring peptides with a positive C-terminal charge. As 111 cross-links were identified with the elastase-trypsin digest, the new peptides with a tryptic C-terminal seemed to have a bigger impact than the increase in complexity. Our analysis so far indicated that reversing the order to trypsin followed by elastase reduces the complexity of the sample and should therefore improve detection of cross-linked peptides. Indeed, more cross-links (131) were identified with the trypsin-elastase digest compared to the elastase-trypsin digest. As a final test of our workflow, we fitted the detected cross-links into the crystal structure of HSA (PDB: 1AO6, monomeric) (Figure 3b). As expected from the link-level FDR (5%), 5% of the links derived from the trypsin-elastase digest were overly long (Figure 3c).

**Figure 3.**
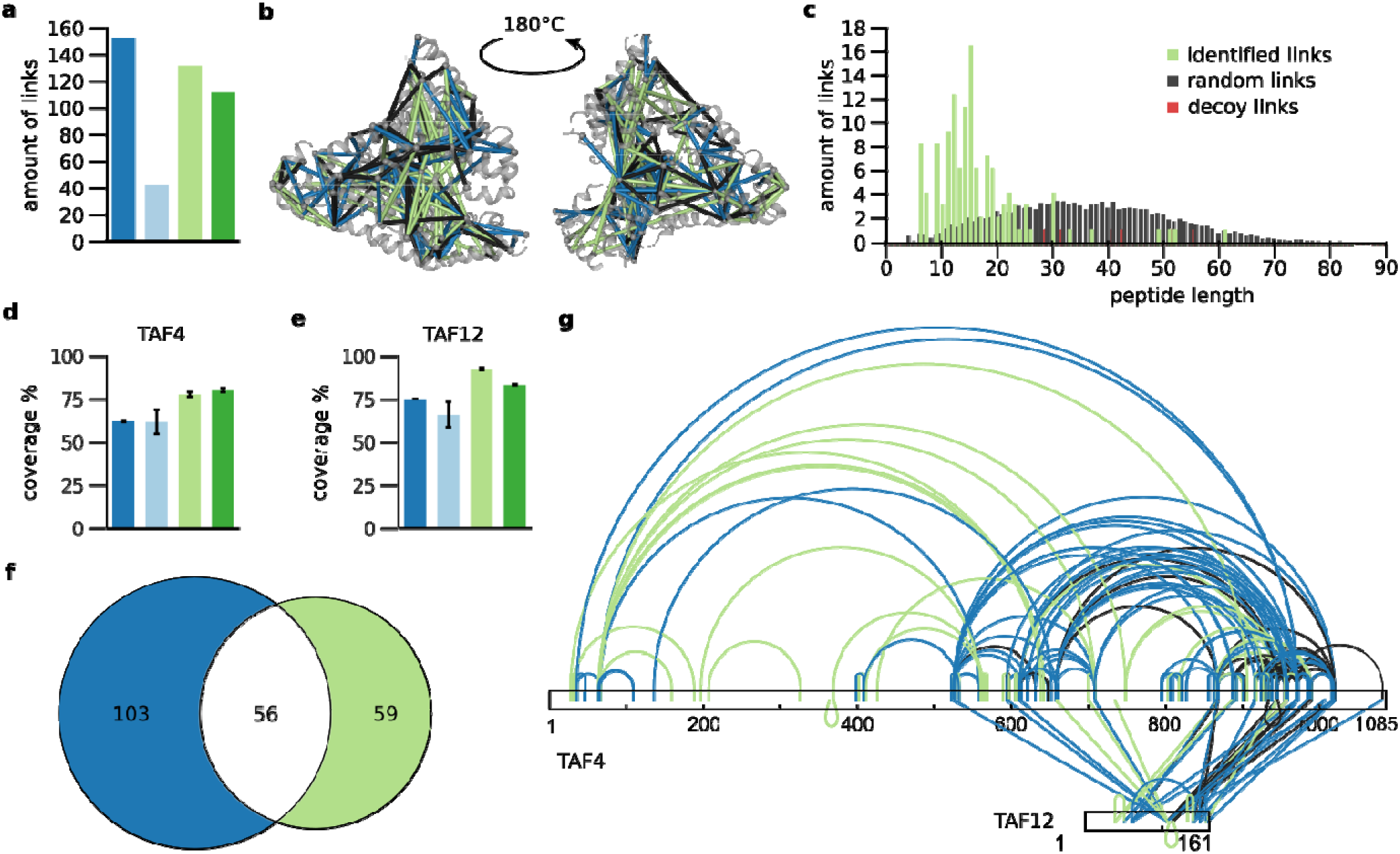
Impact of trypsin-elastase on detection of cross-linked residues. (**a**) Cross-links fitted into the crystal structure of HSA (monomer, PDB: 1AO6) **(b)** Distribution of distances from random links compared to links identified by the sequential trypsin-elastase digest. **(c-d)** Impact of digestion procedure on the sequence coverage of the TAF4-TAF12 complex. Sequence coverage for TAF4 **(c)** and TAF12 **(d)** through linear peptides only. Data shown are the mean ±SD of three independent digestions. **(e)** Venn diagram of identified residue pairs for trypsin and trypsin-elastase sequential digest for the TAF4-TAF12 complex. **(f)** Cross-link map of the TAF4/12 complex for trypsin and trypsin-elastase sequential digest. Trypsin (dark blue), elastase (light blue), sequential trypsin-elastase (light green) and sequential elastase-trypsin digest (dark green).

Having successfully tested our workflow on HSA, we moved on to analyze the TAF4-TAF12 complex. Sequence coverage of both TAF4 and TAF12 decreased when trypsin was replaced by elastase from 74% and 75% to 73% and 65%, respectively (Figure 3d, e). In marked contrast, adding elastase to tryptic peptides increased sequence coverage of TAF4 to 93% and TAF12 to 92%. Reversing the order of digestion to elastase-trypsin still increased the sequence coverage of TAF4 and TAF12 to 95% and 85%. Our previous observation on the impact of complexity and positively charged C-termini on cross-link identification held true for the analysis of the TAF4-12 complex. We identified 162 cross-links with trypsin alone, followed by 115 and 42 cross-links for trypsin-elastase and elastase-trypsin digestion, respectively. A very small number of cross-links (6) was detected when only elastase was used. The cross-links determined from the sequential trypsin-elastase digest were highly complementary to trypsin data with only 59 shared out of 218 identified cross-links (Figure 3f). Importantly, the additional digestion of tryptic peptides by elastase provided data in the previously undetected N-terminal region of TAF4 (Figure 3g).

To test the behavior of elastase in complex mixtures, we applied the protocol to whole cell lysate of *S. pombe*. Three characteristics of the identified peptides – peptide length, peptide type and amount of miscleavages – indicate that the length of the substrate influences the activity of elastase. While digesting tryptic peptides with elastase reduced the number of longer peptides, the median of all digests containing elastase was 13 amino acids (Figure 4a). If the activity of elastase were not influenced by the substrate size, the distribution of tryptic, semi-tryptic and elastase peptides would be independent of the digestion order. However, sequential trypsin-elastase digestion led to more tryptic peptides than elastase-trypsin digestion (Figure 4b). Shorter peptides were very poor substrates for the second enzyme. Consequently, the first enzyme produced the majority of the peptides. Due to the favorable mass spectrometric behavior of tryptic peptides, trypsin should therefore always be the first enzyme used in a sequential digestion. Unsurprisingly, a higher number of miscleavages occurred when elastase was used in all enzyme combinations (Figure 4c). Interestingly, treating the sample with trypsin prior to elastase digestion increased miscleavages. This suggests that the treatment with trypsin reduced the available elastase sites, presumably through their “protection” in smaller peptides.

**Figure 4.**
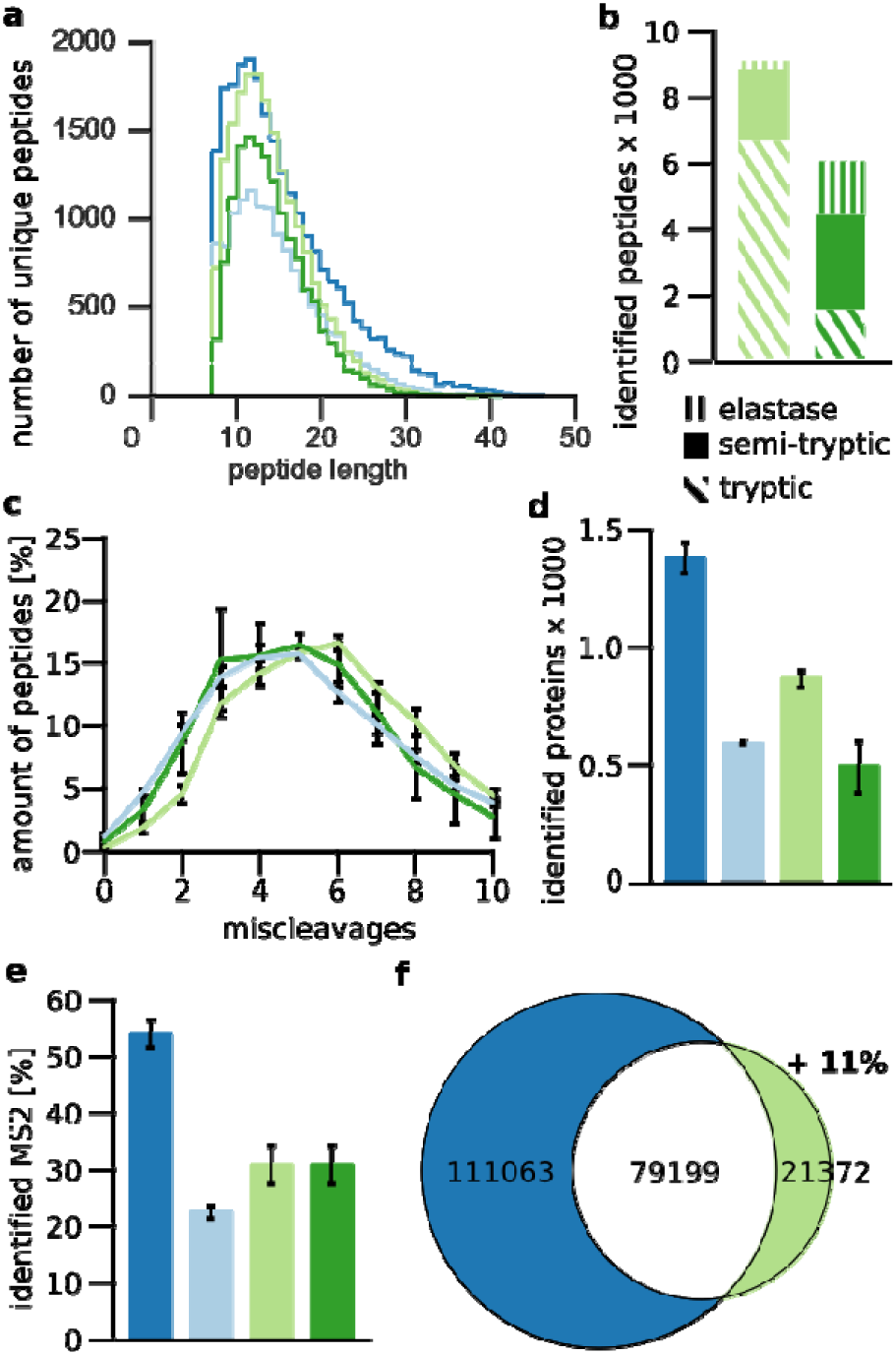
Impact of elastase when following trypsin in a sequential digest in *S. pombe* cell lysate. (**a**) Peptide length distribution. **(b)** Number of tryptic, semi-tryptic and elastase peptides identified in both trypsin-elastase and elastase trypsin digest. (**c**) Distribution of miscleavages for elastase, trypsin-elastase and elastase-trypsin digest. (**d**) Number of identified proteins. **(e)** Percentage of identified MS2. (**f**) Overlap of observed residues by trypsin and sequential trypsin-elastase digestion. Data shown are the mean ± SD of duplicate injections from three independent digestions. Trypsin (dark blue), elastase (light blue) and sequential trypsin-elastase digest (green).

The highest number with 1377 ±62 protein identifications was achieved using only trypsin (Figure 4d). Subsequent elastase digestion reduced the numbers to 874 ±39. Using only elastase yielded 593 ±6 protein identifications. The least amount of proteins, 492 ±115 proteins, was identified with sequential elastase-trypsin. More of the identified semi-tryptic peptides from the sequential digests had a tryptic C-terminus (55 ±2 %), underlining the favorable property of a positively charged residue at the C-terminus for MS/MS-analysis. This may also explain in part the reduced identification numbers in the sequential digestion compared to trypsin alone. Peptides with a non-tryptic C-terminus should be as frequent in the mixture as with a non-tryptic N-terminus. A reduced number of identification paired with a similar number of MS2 spectra between trypsin and sequential digestion, suggests that the challenge is in MS2. Indeed, while the number of MS2 for the trypsin (35203 ±3088) and the sequential trypsin-elastase digest (35160 ±3322) were quite similar, only 31 ±3% from the trypsin-elastase digest compared to 54 ±2% MS2 spectra from the tryptic digest could be matched (Figure 4e). In terms of sequence coverage, the larger number of peptides identified in the trypsin digest also resulted in more residues being covered (190,261 ±8215) than with the sequential digest (100,570 ±10660). However, peptides derived from the sequential digest covered some different regions than tryptic peptides, expanding the covered sequence space of trypsin by 11% (Figure 4f).

## Conclusion

Elastase digestion leading to complementary data compared to trypsin digestion has been shown before and could be confirmed in our study^14,30^. Although sequential digests have been used before^31–33^, to our knowledge elastase has not been exploited in sequential digestion on tryptic peptides or for CLMS. Elastase generates very complex peptide mixtures with many peptides having sequence overlap. Conceptually, CLMS does not combine well with the use of elastase or other proteases with broad cleavage specificity. The increase of complexity and loss in sensitivity normally accompanying the occurrence of overlapping peptides is exacerbated in cross-linking. Every cross-linked peptide consists of two peptides and consequently is subject to combinatorial loss. This meets an already low abundance of cross-linked peptides. The cleavage by trypsin prior to elastase digestion reduces the complexity that is typically associated with elastase digestion. The cleavage of tryptic peptides by elastase is biased towards long peptides and increases their detection. As a result, we detect cross-links in the difficult-to-digest N-terminal region of TAF4. These gains transfer to proteomics at large as shown by our results in *S. pombe* lysate. We anticipate that our protocol will therefore also be useful for other proteomic applications. This includes the detection of post-translational modifications and the analysis of transmembrane domains. The former benefits from increased sequence observation while trypsin leads to systematically large peptides in the latter.

## AUTHOR INFORMATION

### Author Contributions

T.D. and J.R. conceived this study and interpreted data, T.D: conducted all experiments, K.G. and I.B. contributed material, T.D. and J.R. wrote the manuscript with input from all authors. All authors have given approval to the final version of the manuscript.

#### Notes

The authors declare no competing financial interest. The mass spectrometry proteomics data have been deposited to the ProteomeXchange Consortium^34^ via the PRIDE partner repository with the data set identifier PXD011459.

## ACKNOWLEDGMENT

This work was supported by a research stipend to TD [DA 1861/2-1] of the Deutsche Forschungsgemeinschaft, and by the Wellcome Trust through a Senior Research Fellowship to JR [103139], Senior Investigator Award to IB [106115/Z/14/Z] and a multi-user equipment grant [108504]. The Wellcome Centre for Cell Biology is supported by core funding from the Wellcome Trust [203149]. The BrisSynBio Centre is supported by BBSRC/EPSRC [BB/L01386X/1].

---

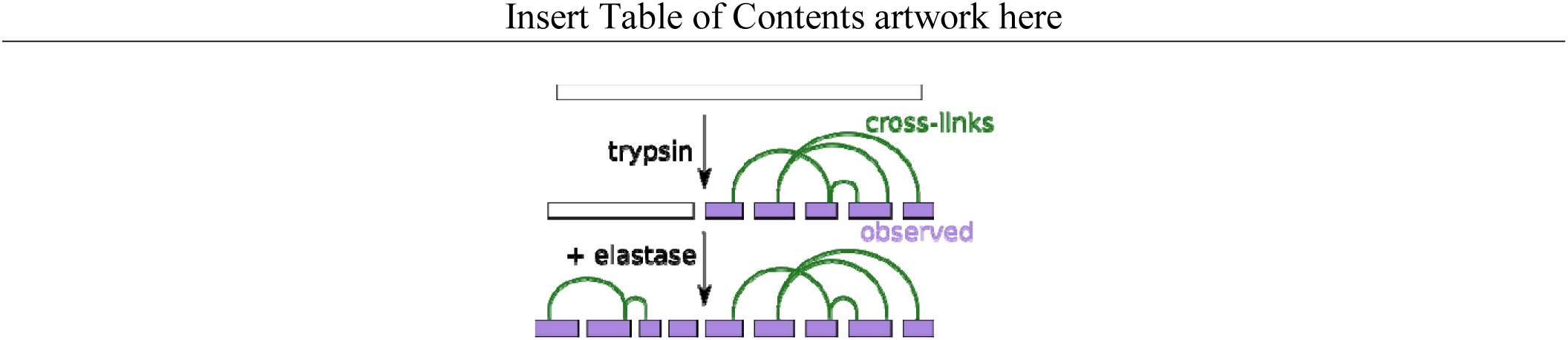

